# Branch-Wise Regularization for Heterogeneous GNN-Based MALDI-TOF AMR Prediction with Biomarker-Consensus Edges

**DOI:** 10.64898/2026.09.24.754085

**Authors:** Wan-Yu Tsai, Yin-Tzer Shih, Y.-H. Chen

## Abstract

This paper predicts antimicrobial resistance (AMR) from matrix-assisted laser desorption/ionization time-of-flight (MALDI-TOF) mass spectra using DRIAMS, evaluated across 13 species-antibiotic datasets spanning sensitive/resistant imbalance ratios of 2.95:1 to 141.21:1 and raw spectral similarities of 0.9166–0.9946. We propose HetSAGE, a heterogeneous graph neural network (GNN) that combines full-spectrum “raw” edges with consensus “biomarker” edges derived from a 4-method feature-selection vote and applies per-edge-type dropout to regularize the two views separately. HetSAGE outperforms a plain multilayer perceptron (MLP) only when spectral similarity is high, consistent with oversmoothing effects reported in the broader GNN literature; the biomarker view converges on features close to independently established clinical markers: within ±1 Da in *Staphylococcus aureus*, Oxacillin, and within range in *Klebsiella pneumoniae*, Meropenem, confirming it is not a black box. At the same time, per-edge-type dropout protects this smaller signal from dilution by regularization tuned for the noisier raw spectrum. Together, these results indicate that heterogeneous graph structure is not a universal win for MALDI-TOF AMR prediction but a conditional one, tied to a measurable property of the data, spectral similarity, rather than to architecture alone. This caveat is largely unaddressed in current DRIAMS-based GNN work and provides a concrete signal for practitioners deciding when a graph model is worth the added complexity over a plain MLP.

**Author Summary:** Diagnosing which antibiotics will work against a bacterial infection typically takes days because it requires growing the bacteria in the lab. A faster technology, MALDI-TOF mass spectrometry, produces a chemical fingerprint of the bacteria within hours, and prior work has shown that this fingerprint alone can predict antibiotic resistance using machine learning. We build on this by connecting samples with similar fingerprints into a graph and using a graph neural network to share information across similar cases. We add a second, more targeted connection type based on a small set of chemical features that a panel of methods agrees are diagnostic, and show that these features align with markers clinicians already use. We find that this graph-based approach helps most when the raw fingerprints are very similar, but offers no advantage otherwise, an important caveat for others building similar systems, since the benefit depends on a measurable property of the data rather than being universal.

## Introduction

In this research, we used the Database of Resistance against Antimicrobials with MALDI-TOF Mass Spectra (DRIAMS) [1] to obtain MALDI-TOF mass spectrometry data for over 300,000 clinical bacterial isolates from four laboratories in Switzerland. We investigated antimicrobial resistance (AMR) prediction across a panel of 13 species-antibiotic datasets spanning three clinically important pathogens: *Escherichia coli, Klebsiella pneumoniae*, and *Staphylococcus aureus* (see Table 1), using DRIAMS-A, the largest sub-collection. We developed a heterogeneous graph neural network from these data to predict antibiotic resistance across this panel. Fig 1 provides an overview of this end-to-end process, from data collection through feature engineering, model development, and evaluation.

**Table 1.** Study Panel Characteristics: Class Imbalance and Spectral Similarity Across 13 Species-Antibiotic Datasets from DRIAMS-A.

| Species | Antibiotic | S/R ratio | sim<sub>raw</sub> | n.sample |
| --- | --- | --- | --- | --- |
| *E.coli* | Ciprofloxacin | 2.95 | 0.9931 | 7,885 |
| *E.coli* | Ceftriaxone | 4.47 | 0.9892 | 8,112 |
| *E.coli* | Cefepime | 11.54 | 0.9865 | 7,159 |
| *E.coli* | Piperacillin-T. | 14.46 | 0.9914 | 7,576 |
| *E.coli* | Tobramycin | 7.03 | 0.9946 | 4,876 |
| *K.pneumoniae* | Ciprofloxacin | 7.31 | 0.9827 | 5,454 |
| *K.pneumoniae* | Ceftriaxone | 7.61 | 0.9412 | 5,623 |
| *K.pneumoniae* | Cefepime | 19.56 | 0.9166 | 5,120 |
| *K.pneumoniae* | Meropenem | **141.21** | 0.9166 | 3,413 |
| *K.pneumoniae* | Tobramycin | 7.81 | 0.9826 | 2,953 |
| *S.aureus* | Ciprofloxacin | 8.83 | 0.9811 | 6,922 |
| *S.aureus* | Fusidic acid | 15.48 | **0.9942** | 4,795 |
| *S.aureus* | Oxacillin | 5.19 | 0.9796 | 4,905 |

**Fig 1.**
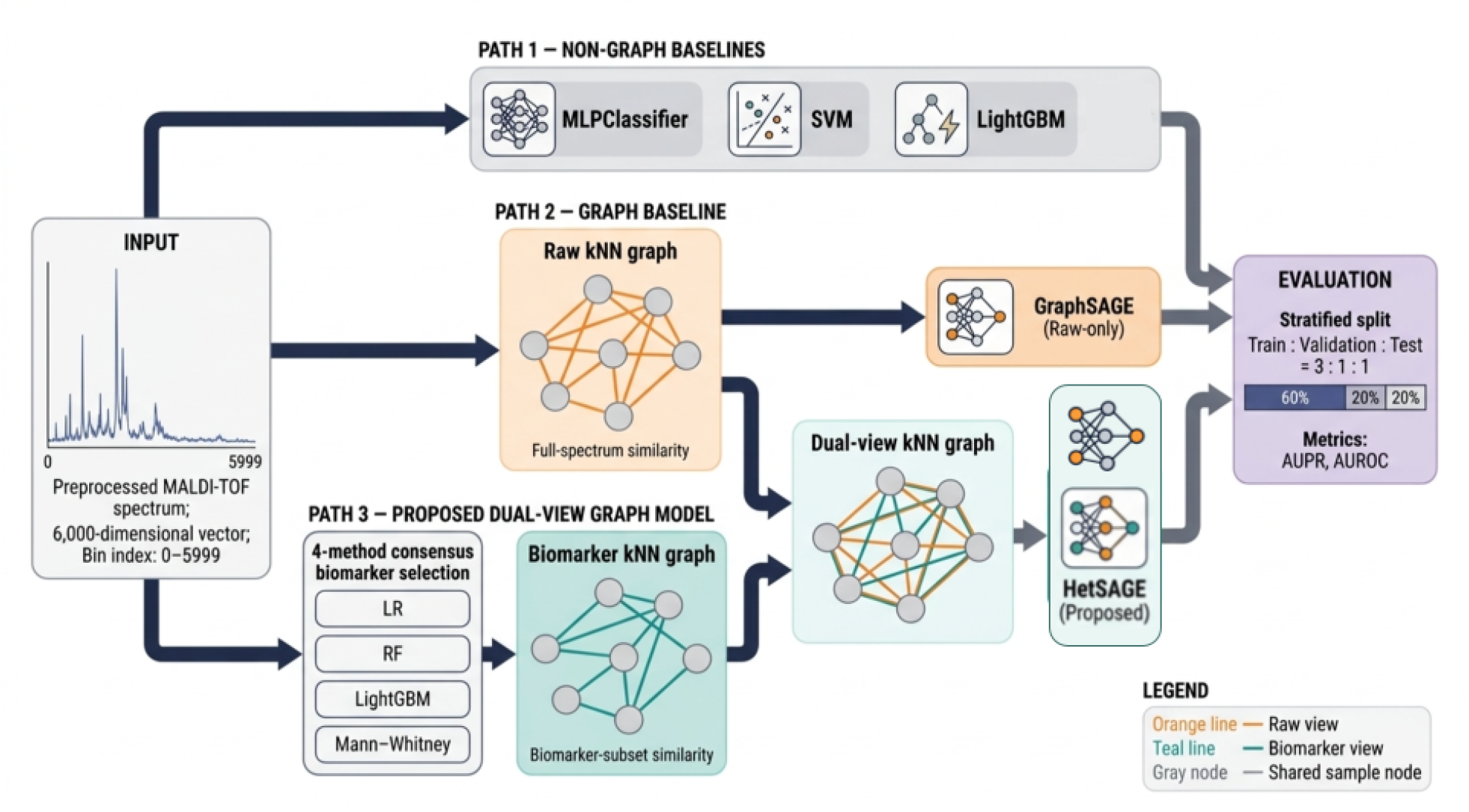
Overview of the Proposed Approach: Data Collection to Model Evaluation.

Traditional culture-based phenotypic resistance testing can take up to 72 hours from sample collection to a reported result: a figure corroborated across independent sources, reported as “up to 72 hours” [1] and as a “48–72 hours” range [2]. The delay is driven largely by the 24–48 hours required for bacterial growth [3]. MALDI-TOF mass spectra, by contrast, are routinely acquired within 24 hours of sample collection for species identification [1–3]: the same spectrum that traditional workflows then hand off to a separate, slower phenotypic antibiotic susceptibility testing (AST) step to determine resistance. Weis et al. [1], who created DRIAMS, showed that this already-acquired spectrum is sufficient on its own. They trained calibrated classifiers such as logistic regression, LightGBM, and a multilayer perceptron directly on the MALDI-TOF spectrum to predict antimicrobial resistance, without requiring genomic sequencing or waiting for the additional phenotypic AST step, bringing the total time to a resistance result down to roughly 24 hours, the time of the initial MALDI-TOF step itself (see Fig 2). Validated on a panel of clinically important pathogens, their models reached an area under the ROC curve (AUROC) of 0.80 and an area under the precision-recall curve (AUPR) of 0.49 for oxacillin resistance in *S. aureus*, and an AUROC of 0.74 for ceftriaxone resistance in both *E. coli* (AUPR 0.30) and *K. pneumoniae* (AUPR 0.33); LightGBM performed best for *E. coli* and *S. aureus*, while the multilayer perceptron performed best for *K. pneumoniae*. A retrospective clinical case study of 63 patients also found that this approach would have changed treatment in nine cases, benefiting eight (89%) [1]. This showed that MALDI-TOF spectra alone provide sufficient discriminatory signal to predict resistance in roughly a third of the traditional turnaround time for the pathogens in this study’s panel.

**Fig 2.**
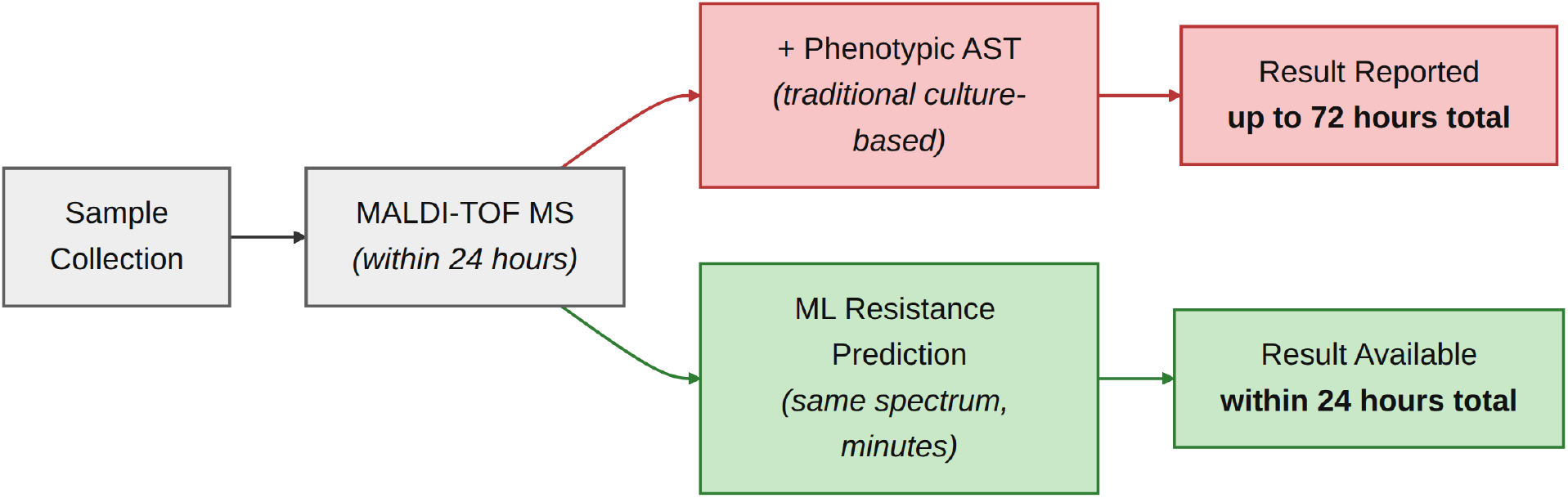
Turnaround Time Comparison: Traditional Phenotypic AST Versus MALDI-TOF Machine Learning Prediction.

We aimed to develop a rapid identification model to assess the potential effectiveness of these antibiotics against the corresponding pathogens. Using our machine learning models, we can quickly determine antibiotic resistance in cultured clinical bacterial isolates after MALDI-TOF analysis. This method greatly shortens the time required compared with conventional antibiotic susceptibility tests. We detail the experimental setup, cross-validation protocol, and evaluation metrics in Experiments and Results.

## Related Work

Several recent studies build directly on DRIAMS or use it alongside other public MALDI-TOF repositories. Wang et al. [4] trained MALDI-TOF-plus-ML classifiers for *S. aureus* mecA and AMR status using both DRIAMS and an independent Taiwan hospital database, the closest existing precedent to a cross-population framing of this panel. Wang et al. [5] built a deep learning system that identifies bacterial species, predicts resistance, and recommends antibiotics directly from raw MALDI-TOF spectra, training on DRIAMS (2015–2018) together with 89,026 local hospital records: the same research group that later reported multicenter AUROCs up to 0.95 in [2]. Astudillo et al. [6] took a multi-label approach, predicting resistance to several antibiotics jointly from a single model across multiple DRIAMS sub-collections, in contrast to the per-species-antibiotic binary panel used here. Chen et al. [7] benchmarked cross-site generalization directly, evaluating whether models trained at one hospital transfer to another; their central finding, that a modest amount of target-site labeled fine-tuning closes most of the domain gap, while unsupervised domain-adaptation methods add little, is directly relevant to any claim that this panel’s models would generalize beyond DRIAMS-A. Dai et al. [8] proposed a region-guided data-augmentation method trained on a private *E. coli* levofloxacin dataset and validated it by transferring it to DRIAMS-C ceftriaxone, relevant to this panel’s *E. coli* subset. Wiesmann et al. [9] is not a DRIAMS study but an independent validation: they built their own German MALDI-TOF-plus-AMR database, structured analogously to DRIAMS-A, and found that models trained at one site still degrade without adaptation when applied elsewhere, corroborating the same cross-site limitation that Chen et al. [7] benchmark directly. Corvelo Benz et al. [10] also train directly on DRIAMS-A, but target a different problem than raw prediction accuracy: a knowledge-graph-enhanced conformal predictor that wraps a GNN-based classifier (ResMLP-GNN) to issue statistically guaranteed, multi-antibiotic resistance prediction sets for *K. pneumoniae*, trading the single-antibiotic binary framing used here for simultaneous coverage guarantees across an antibiotic panel.

Chen et al. [7] is also the only surveyed work to combine DRIAMS [1] with two other public repositories, MS-UMG (Germany) and MARISMa (Spain, different instrument), benchmarking 13 methods (3 baselines, 4 supervised transfer methods including Fine-Tuning, the best performer, and 6 unsupervised domain-adaptation (DA) methods, all sharing one MLP backbone) and finding that fine-tuning with 25% of target-site labels recovers 81–94% of full performance, versus only 0–6% for unsupervised DA. Among the 13, GNN-DA and GNN-DA-FT use a graph neural network over a drug co-resistance graph whose nodes are species-antibiotic pairs, applied to cross-site domain adaptation, unlike Method, which builds a per-sample graph with “raw” and “biomarker” edge types for within-site prediction. Both MS-UMG [11] and MARISMa [12] are openly downloadable via Zenodo with no application needed; MS-UMG (CC0, 30.6 GB, pre-binned to 6,000-dim) has been downloaded and verified locally, with cross-site experiments extending Method’s approach to it planned as future work, while MARISMa (CC-BY-4.0, 16.8 GB raw spectra + AMR.csv) ships unbinned and was not downloaded.

### Graph Neural Network Architectures

The Graph Convolutional Network (GCN) [13] aggregates neighbor features using a fixed, degree-normalized average and a single shared weight matrix per layer on a single homogeneous graph. The Graph Attention Network (GAT) [14] replaces the fixed average with a learned attention weight per edge, allowing some neighbors to matter more than others, still on a single edge type. GraphSAGE (Graph SAmple and aggreGatE) [15] instead samples a fixed-size neighborhood and learns an aggregator (mean, LSTM, or pooling) that generalizes to unseen nodes at inference time: the inductive property Method relies on, and the base aggregator each of HetSAGE’s two branches uses. None of these three handle more than one edge type per graph. The Heterogeneous Graph Neural Network (HetGNN) [16] and the Relational Graph Convolutional Network (R-GCN) [17] generalize to heterogeneous graphs by giving each relation or node type its own transformation before aggregation. HetGNN does this via type-specific neighbor sampling and encoders, and R-GCN via a per-relation weight matrix, while the Heterogeneous graph Attention Network (HAN) [18] instead defines heterogeneity through hand-picked meta-paths combined via a learned attention. Method’s HetSAGE follows the same type-specific-branch principle as HetGNN and R-GCN, but with only two fixed, domain-defined edge types (raw, biomarker) rather than an arbitrary relation set, and combines them via semantic attention rather than meta-paths.

In biology, GNNs are most established in domains where a graph is already native to the data: molecular property and drug-target/drug-drug interaction prediction, where atoms and bonds form the graph directly [19]; protein structure prediction, where AlphaFold’s Evoformer applies attention over residue pairs [20]; and single-cell omics, where GNNs model cells or genes as nodes over epigenomic, transcriptomic, spatial, and multi-omics graphs [21]. This is the same omics setting already flagged in Dataset [22, 23] as one where GNNs often fail to beat simpler baselines. MALDI-TOF AMR prediction has no native graph at all: spectra are binned intensity vectors, not molecules or cells; so Method’s raw/biomarker *k*NN graph is a constructed rather than a given structure, closer in spirit to the omics setting’s mixed track record than to the domains where GNNs are best established. Constructing a *k*NN or similarity graph over samples, rather than using a given molecular or interaction graph, has precedent in biology: Parisot et al. [24] build a sparse population graph over patients from imaging-feature similarity for GCN-based Autism/Alzheimer’s classification, and Zeng et al. [25] build a spot-level *k*NN graph from Euclidean distance, feeding it into a GraphSAGE-based GNN for spatial transcriptomics prediction. This is the same *k*NN-then-SAGE pattern Method uses, though over spots rather than per-sample spectra.

Nguyen et al.’s AMR-GNN [26] is a closer methodological precedent still, applying a dual-GCN, multi-representation architecture (distance-thresholded *k*NN graphs over unitig, single nucleotide polymorphism (SNP), and frequency chaos game representation (FCGR) genomic features, fused via low-rank multimodal fusion) to AMR prediction directly, and reporting very major error (VME) and major error (ME) rates alongside AUROC as clinically relevant metrics. This is the same VME/ME framing motivating Evaluation Metrics’ threshold-metric discussion. Unlike Method’s two fixed, domain-defined edge types (raw, biomarker) over MALDI-TOF spectra, AMR-GNN’s graphs are built from whole-genome-sequencing-derived features, and their edge-count threshold is tuned per representation, with a separate multi-locus sequence typing (MLST)-based decoupling step added afterward to suppress edges between clonally related isolates. AMR-GNN also selects a single best-performing representation as its primary node-feature input before graph construction, unitig, chosen empirically by comparing held-out AUROC across SNP, unitig, and FCGR; Method instead keeps both views (raw, biomarker) permanently and combines them via semantic attention rather than choosing one.

### Class Imbalance Across Domains and Model Capacity

Class imbalance severity is conventionally described in three broad tiers, though the domain determines how far the top tier extends. Existing surveys commonly treat ratios up to roughly 10:1 as mild and label a dataset severely imbalanced once it passes a fixed threshold, typically 50:1 or 100:1 [46]. General-purpose benchmark work concentrates in the 1:4 to 1:100 range, while specific rare-event domains such as fraud detection and cheminformatics treat 1:1000 to 1:5000 as their normal operating range [47]. This panel’s worst-case ratio, 141.21:1 (see Table 1), falls just past the general-purpose severity threshold but well short of the fraud-detection/cheminformatics tail, consistent with describing it as severely, rather than extremely, imbalanced.

Architecture choice under imbalance is better predicted by data structure and sample size than by imbalance ratio alone. Ghosh et al. [48] find that deep multilayer perceptrons are affected by class imbalance to about the same degree as shallow classifiers, so depth alone does not confer robustness to imbalance. A sample-size effect is better established instead: Grinsztajn et al. [49] show tree-based models such as gradient boosting and random forests outperform deep learning across a broad range of tabular dataset sizes, with the gap narrowing only as datasets grow into the tens of thousands of rows or more.

Convolutional neural networks (CNN) and graph convolutional networks (GCN) are rarely compared head-to-head for a related reason: CNNs operate on grid-structured data and GCNs on graph-structured data, so the choice between them follows from whether the data is natively a grid or a graph, not from imbalance ratio.

### Dataset

We use DRIAMS, a publicly available database of MALDI-TOF mass spectra linked to phenotypic antibiotic-resistance labels, collected from four Swiss diagnostic laboratories between 2016 and 2018 [1]. This study uses DRIAMS-A (University Hospital Basel), the largest sub-collection, with 145,341 spectra. Each spectrum is preprocessed into a 6,000-dimensional vector (2,000–20,000 Da, 3 Da bins) [1], which is DRIAMS’s untargeted acquisition range.

Our panel consists of 13 Species × Antibiotic datasets shown in Table 1. S/R imbalance ranges from 2.95:1 to 141.21:1 across the panel; raw spectral similarity (sim_raw_) ranges from 0.9166 to 0.9946. Four datasets, *E*.*coli* Ciprofloxacin, Piperacillin-T., and Tobramycin, plus *S*.*aureus* Fusidic acid, exceed the sim_raw_ *>* 0.99 threshold discussed below: these are the cases where the HetSAGE biomarker edge is expected to matter most; the remaining nine fall in the 0.9166–0.9892 range, where a plain MLP baseline is expected to be more competitive.

High spectral similarity limits how much of an advantage HetSAGE can offer over a plain MLP. We initially proposed HetSAGE, a heterogeneous GNN combining raw full-spectrum *k*NN edges with biomarker consensus *k*NN edges, with *k* selected per dataset according to sim_raw_, to counter two properties of this panel: high inter-sample spectral similarity and severe class imbalance, which degrade prediction quality in the raw 6,000-dim feature space. However, evaluation shows this benefit is conditional, not universal (see Table 5 in Discussion for the full condition-by-condition breakdown). Mechanistically, this conditionality aligns with the broader GNN literature on oversmoothing: when the spectral node features are already highly similar, message-passing aggregation over neighbors compresses representations further rather than adding discriminative signal, reducing or eliminating the graph’s advantage over a model that uses the raw features directly [27]. This is not just a general-GNN-theory concern: it is directly observed in biological/omics prediction tasks structurally similar to ours. A peer-reviewed benchmark on transcriptomic phenotype prediction found GNNs “rarely provide a real improvement” over simpler methods, performing comparably to MLPs, and attributed this partly to the limited predictive quality of the underlying biological network itself [22]. A 2026 benchmarking framework for graph-level prediction in the low-sample, high-dimension omics regime, a setting structurally similar to our 6,000-dim, few-thousand-sample panel, similarly found that “widely used GNNs often do not outperform simple MLPs and classical baselines” [23]. This motivates going beyond the single additional edge type introduced in Method, rather than treating heterogeneity alone as sufficient.

## Method

This section proceeds in three steps. Consensus-Vote Biomarker Construction builds a second, more informative feature view, the biomarker consensus-vote feature set, motivated by the raw spectrum’s noise and high dimensionality, and validates that this second view is not a black box by comparing it with independently established clinical biomarkers. HetSAGE Architecture then defines HetSAGE, the heterogeneous GNN that consumes both views: it builds up a plain GNN, then a heterogeneous GNN, then the specific *k*NN-graph workflow used here, and presents the architecture’s formulas. Per-Edge-Type Dropout uses Consensus-Vote Biomarker Construction’s validation as the basis for treating the two views asymmetrically, giving each its own dropout rate rather than sharing one. Empirical results for this panel are reported separately in Experiments and Results.

### Consensus-Vote Biomarker Construction

The raw spectrum, a noisy 6,000-dimensional input, has already been shown in Dataset that, at high inter-sample similarity, this raw view provides limited information for a plain MLP. Instead of supplying a GNN aggregator with only this noisy data, we create a second, more informative feature view: biomarker. This biomarker combines a 4-method consensus vote to highlight the wavelengths most discriminative of resistance. The features it converges on align with what clinicians already look at. We deliberately chose the four methods to be complementary rather than redundant, each capturing a different kind of signal.

Logistic Regression (LR) coefficient magnitude captures features useful under linear separation. Random Forest (RF) feature importance [28] captures features useful across bagged random subsets. LightGBM (LGBM) feature importance [29] captures features that reduce error under boosting. The Mann-Whitney U test [30] is a model-free statistical check for features where the S and R distributions simply differ. A feature enters the biomarker set only when enough of these four agree, filtering out artifacts specific to any one model rather than trusting a single method’s blind spots. This ensembling-over-a-single-selector approach follows established ensemble feature selection methodology, which shows that combining heterogeneous selectors (linear, bagging, boosting, statistical) is more stable and less prone to any one method’s failure mode than relying on a single method [31]. Support Vector Machine (SVM), used elsewhere in this study as a classification baseline, is deliberately excluded from the vote. A non-linear-kernel SVM has no native per-feature importance score, since the kernel trick maps inputs into an implicit space and the decision function can’t be cleanly decomposed back out of it, documented as a general black-box limitation of non-linear SVMs, not specific to our data [32]. Extracting one would require a dedicated post hoc approximation, reintroducing the extra cost and circularity the consensus vote was designed to avoid; we chose Mann-Whitney instead as the fourth, approximation-free view. In *S*.*aureus*, the consensus feature landed at m/z 2,414, within *±*1 Da of the clinically validated PSM-mec marker at 2,415 *±* 2 Da [33, 34]. In *K*.*pneumoniae*, the consensus feature at m/z 11,399 fell within the range of the Bruker-validated p019 marker at 11,109 Da: the same marker already used in automated KPC detection at 97% sensitivity [35–37]. That’s genuine biological validation (Tier 1, established causal chain), not just agreement between two ML studies (Tier 2): the LGBM-based marker reproductions shown in Fig 5 and Fig 6 (see S1 Appendix) [38, 39]. So the biomarker is not a black box; it recovers signal that independently derived clinical assays also rely on, giving the GNN aggregator defined in HetSAGE Architecture a genuinely helpful second input rather than a second noisy one.

#### Algorithm 1

GraphSAGE Forward Pass (Single Edge Type *t, k*-Neighborhood).

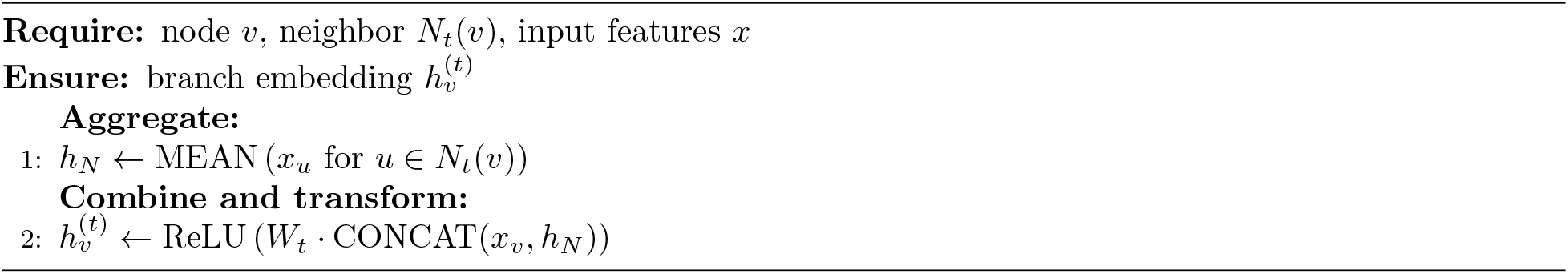

### HetSAGE Architecture

#### Graph construction

A plain GNN passes each node’s features through layers that aggregate information from its neighbors, letting a node’s representation be shaped by its connections rather than by its own features alone [13], but this requires a graph to already exist. MALDI-TOF spectra have no native edges: DRIAMS provides isolate-level spectra and resistance labels, not any given link between isolates.

The graph must therefore be constructed before any GNN layer can run. We do this with *k*NN: each sample becomes a node, and for each edge type *t ∈ {*raw, biomarker*}*, a node *v* is linked to its *k* nearest neighbors *N*_*t*_(*v*) under *t* ’s own similarity metric: full-spectrum similarity for the raw edge type, or consensus similarity, defined in Consensus-Vote Biomarker Construction, for the biomarker edge type. *k* is selected per dataset according to sim_raw_. A heterogeneous GNN extends this idea to graphs with more than one relation type, giving each relation or edge type its own aggregation before combining the results [16, 17], as reviewed in Graph Neural Network Architectures: here, that means building two separate *k*NN graphs over the same samples, one per edge type, rather than one graph with mixed edges.

#### Notation

Table 2 summarizes the symbols used in this section and Per-Edge-Type Dropout.

**Table 2.** Notation Used in HetSAGE Architecture and Per-Edge-Type Dropout.

| Symbol | Meaning |
| --- | --- |
| $v$ | a node (sample) |
| $t \in \{\text{raw}, \text{biomarker}\}$ | edge type / branch |
| $N_t(v)$ | node $v$'s $k$ nearest neighbors under edge type $t$ |
| $x_v$ | node $v$'s input feature vector |
| $x_v^{(t)}$ | $x_v$ after branch-$t$ dropout |
| $W_t$ | branch-$t$ weight matrix (GraphSAGE transform) |
| $h_v^{(t)}$ | branch-$t$ embedding for node $v$ |
| $h_v$ | combined (post-attention) embedding for node $v$ |
| $p_{\text{raw}}, p_{\text{biomarker}}$ | per-branch dropout rates |
| $\alpha_v^{(t)}$ | semantic-attention weight for branch $t$ at node $v$ |
| $W_a, b_a, q$ | semantic-attention parameters |
| $y_v$ | predicted resistance probability for node $v$ |

#### Encoding

Once the two *k*NN graphs exist, HetSAGE encodes each edge type through its own branch, using the standard GraphSAGE mean-aggregator forward pass [15] unmodified, run once per edge type *t* ∈ {raw, biomarker}, each with its own weight matrix. Algorithm 1 details this forward pass for a single edge type:

*x*_*v*_ is node *v*’s input feature vector: the full 6,000-dim spectrum for the raw branch, or the biomarker consensus feature subset from Consensus-Vote Biomarker Construction for the biomarker branch. *W*_*t*_ is a branch-specific weight matrix. This is unmodified, textbook GraphSAGE: nothing about it is specific to this panel or to heterogeneity yet. Per-Edge-Type Dropout explains how the two resulting branch embeddings, 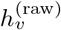 and 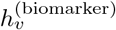, are combined into HetSAGE and presents the one change we propose to this step.

##### Algorithm 2

HetSAGE Forward Pass (Edge Types *t ∈ {*raw, biomarker*}*).

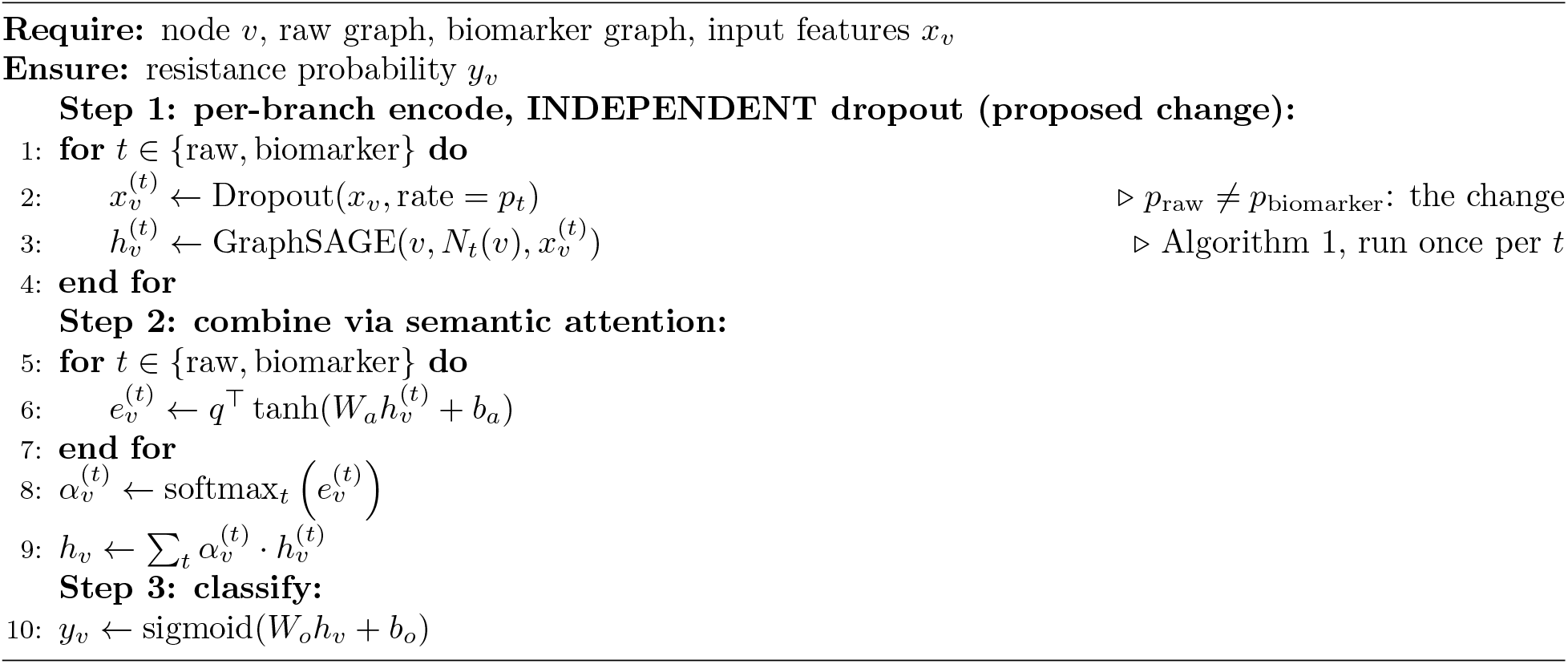

### Per-Edge-Type Dropout

Fig 3 summarizes this end-to-end: each sample’s raw and biomarker views enter separate GraphSAGE encoder branches (see HetSAGE Architecture), producing 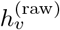 and 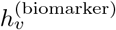; the two branch embeddings are then combined using a learned, per-sample semantic-attention weight rather than a fixed average into a single embedding *h*_*v*_, which a final linear-plus-sigmoid layer maps to a resistance probability *y*_*v*_. Algorithm 2 presents HetSAGE’s full forward pass and highlights this paper’s proposed change:

**Fig 3.**
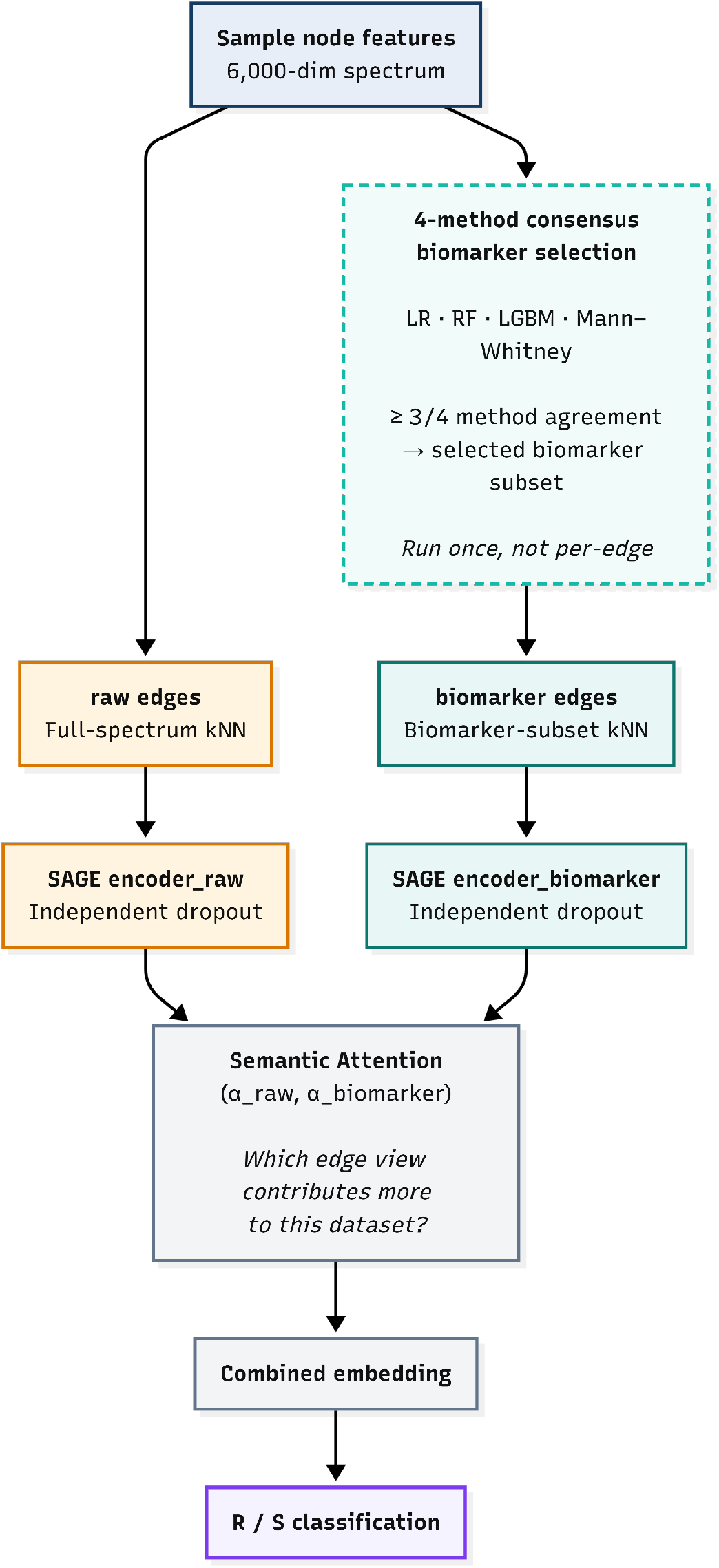
HetSAGE Architecture and Workflow: Combining Raw and Biomarker Views.

Prior to this proposal, Step 1 used a single shared dropout rate *p* for both branches. raw is the noisy, 6,000-dimensional full spectrum; it can absorb fairly aggressive regularization without losing much. biomarker, by contrast, is already the distilled, clinically-validated signal established in Consensus-Vote Biomarker Construction, applying the same dropout rate to it risks discarding exactly the bits just confirmed to matter. Splitting *p* into *p*_raw_ ≠ *p*_biomarker_ costs nothing architecturally; it only means separating one hyperparameter into two, one per branch: exactly the two encoder boxes Fig 3 shows in parallel. This is most consequential exactly where Dataset showed the panel struggles: the high-similarity, high-dimensional datasets, where raw alone is least informative and biomarkers’ smaller, validated signal most needs protecting rather than diluting. Experiments and Results reports how this plays out empirically across the panel.

## Experiments and Results

### Experimental Setup

During data preprocessing, mass spectrometry data were segmented into fixed intervals from 2,000 Da to 20,000 Da in 3-Da intervals, producing a 6,000-dimensional vector for each sample (see Dataset). We created binary labels: sensitive (S) or resistant (R), based on the antibiotic resistance categories. To predict AMR across the panel in Table 1, we evaluated five models: HetSAGE, the proposed heterogeneous GNN with raw+biomarker edges (see Method); GraphSAGE; MLPClassifier; SVM; and LightGBM (LGBM). This design isolates the contribution of heterogeneity by comparing HetSAGE against GraphSAGE, and the contribution of using a graph at all by comparing either GNN against the three non-graph baselines.

Fig 4 summarizes this end-to-end experimental pipeline. After preprocessing, the panel splits into three paths, comparing three groups of models: a no-graph path that goes directly to the three non-graph baselines (MLPClassifier, SVM, LGBM); a raw-only path through a single *k*NN graph into GraphSAGE, the homogeneous GNN (Algorithm 1); and the combined raw+biomarker path, pairing the full-spectrum view with the biomarker view from Consensus-Vote Biomarker Construction, into HetSAGE, the proposed heterogeneous model with per-edge-type dropout, whose internal architecture is detailed separately in

**Fig 4.**
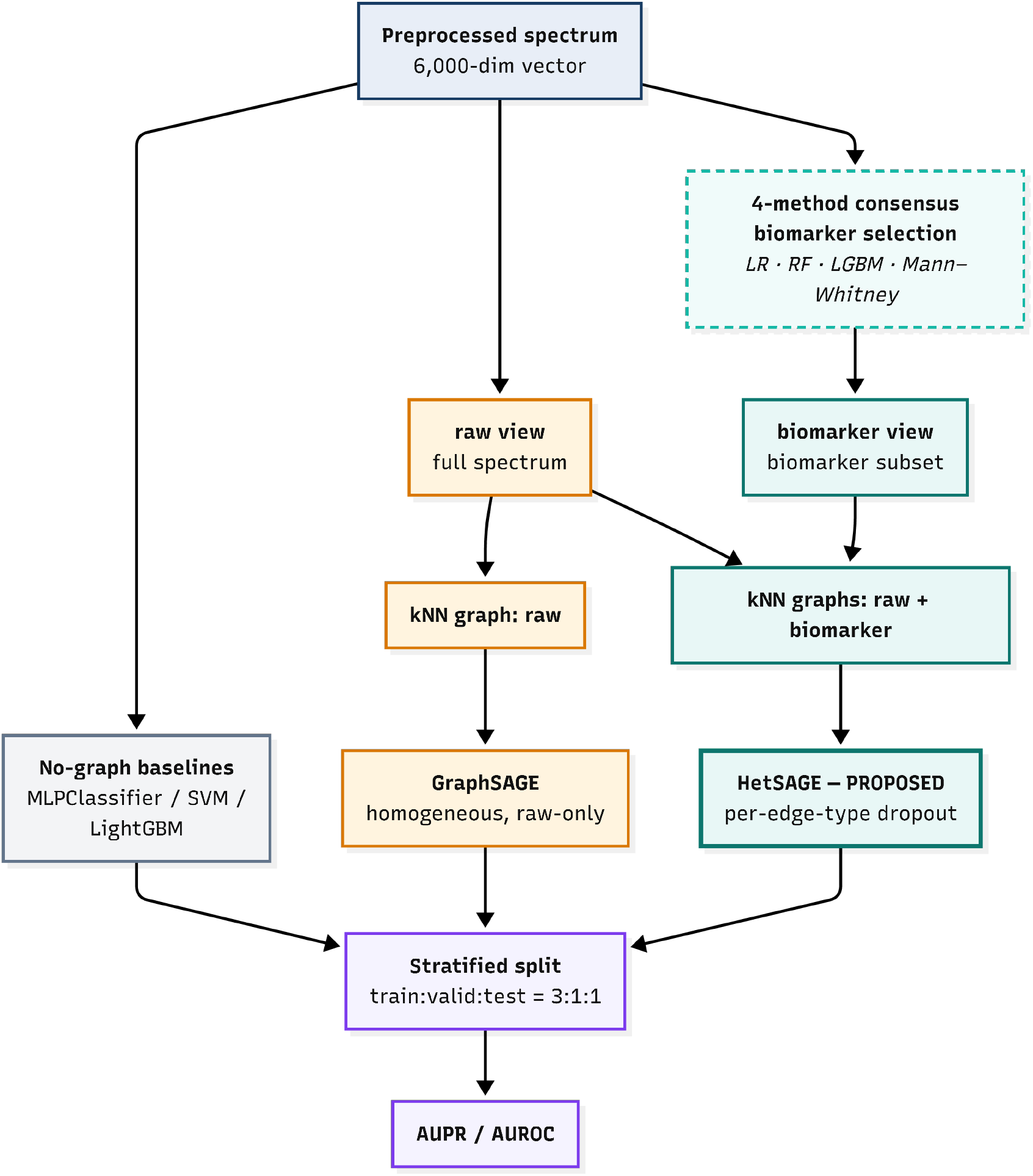
Experimental Pipeline: Three Model Paths from Preprocessing to Evaluation.

**Fig 5.**
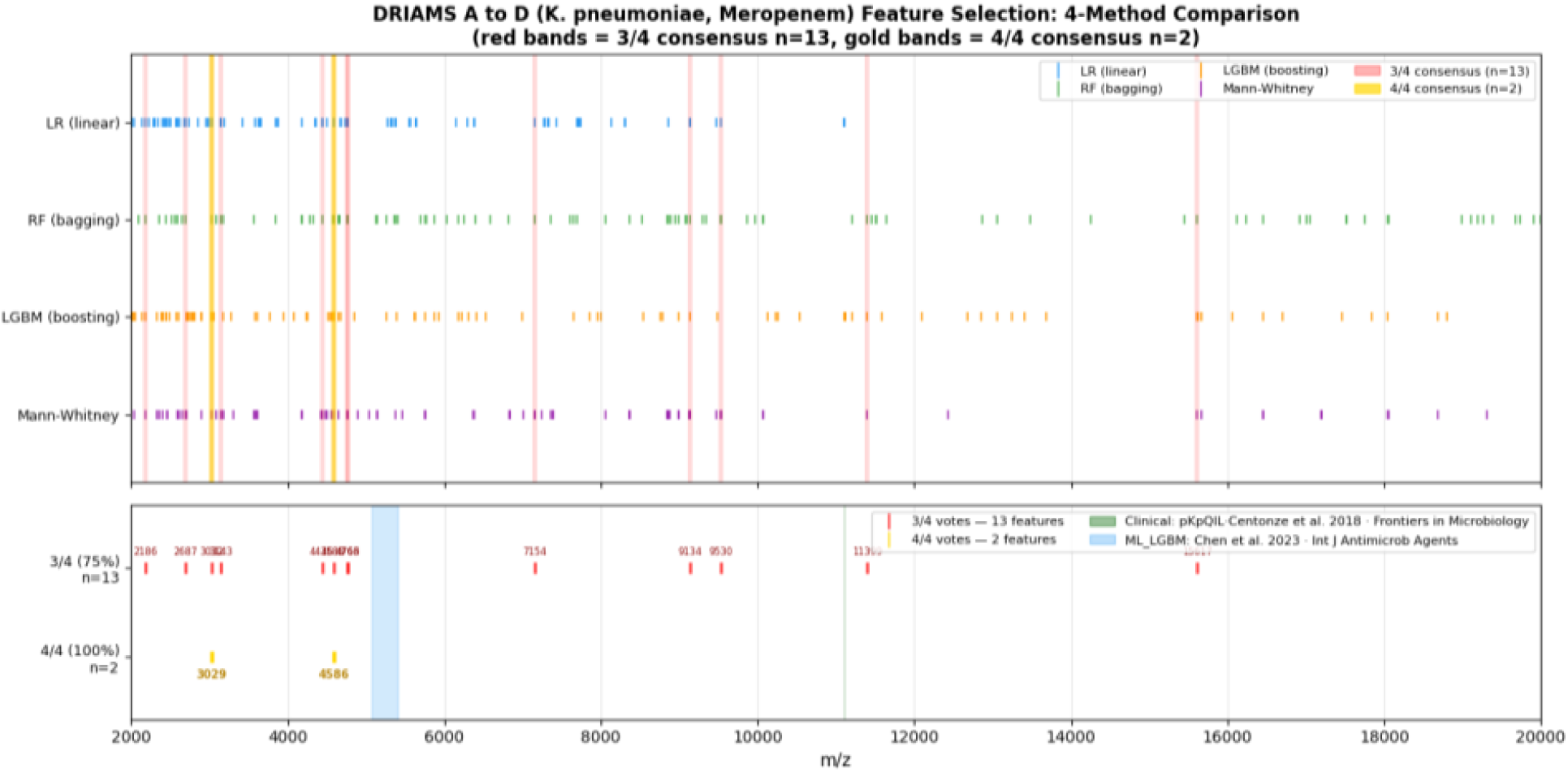
*K*.*pneumoniae*, Meropenem: 4-Method Feature Selection Comparison.

**Fig 6.**
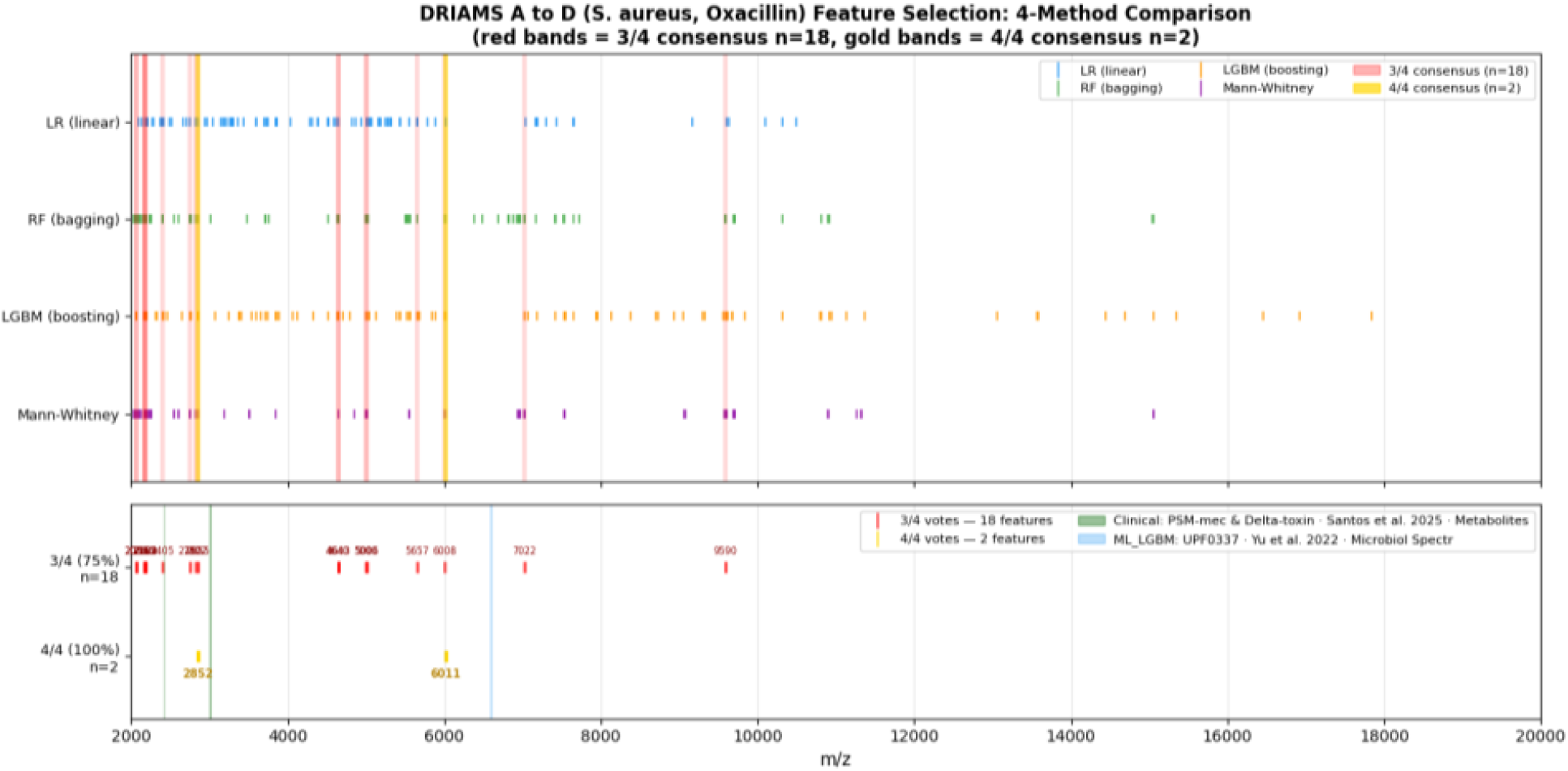
*S*.*aureus*, Oxacillin: 4-Method Feature Selection Comparison.

Fig 3 (see Per-Edge-Type Dropout). All three paths converge on the same stratified train:valid:test split (see Cross-Validation Protocol) and on AUPR/AUROC evaluation (see Evaluation Metrics and Results).

### Cross-Validation Protocol

We used stratified 4-fold cross-validation rather than a single held-out split and reported mean ± std across folds for all five models on all 13 datasets in Table 1. Stratification preserves each fold’s S/R class ratio at the dataset’s own imbalance level (up to 141.21:1; see Table 1), rather than letting a random split starve a fold of already-scarce resistant (R) samples.

Concretely, each dataset is first stratified-split into a fixed 20% holdout test set, set aside from all training and model selection, and scored only once, at the end. The remaining 80% (the learning set) is itself stratified into 4 folds; across the 4 rotations that give this protocol its name, 3 folds serve as training data, and 1 serves as validation for model selection, while the same fixed test partition is evaluated in every rotation. This gives each rotation a train:validation:test ratio of 3:1:1 over the original five-way partition. This follows the standard hold-out-plus-k-fold pattern for keeping the final test set fully unseen during model selection: the test partition is never trained or validated on at any point, unlike a plain single train/valid/test split where the same held-out data risks being reused across tuning decisions.

### Evaluation Metrics

Both AUROC and AUPR are reported below, but AUPR is treated as the more decision-relevant metric for this panel because of how the two curves respond to class imbalance, not preference alone. Let *P* and *N* denote the number of positive (R) and negative (S) samples, with *N≫ P* (i.e., *S ≫ R*) under the severe imbalance seen in this panel (up to 141.21:1; see Table 1). AUROC and AUPR are both computed from the same confusion-matrix counts, but they normalize false positives (*FP*) against different denominators:

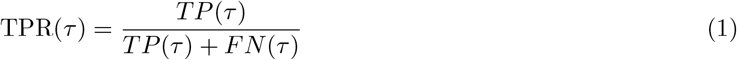

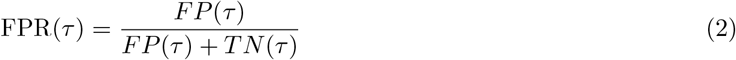

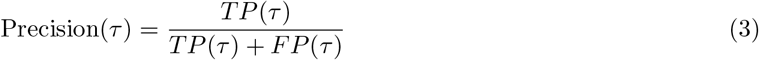

These are the standard confusion-matrix-derived rates [40, 41], each evaluated at a specific decision threshold *τ* : the underlying counts *TP, FP, TN, FN* change as *τ* varies, so TPR, FPR, and Precision each take a different value at each *τ* [40]. Sweeping the decision threshold *τ* across its full range traces out two curves from these per-threshold quantities, and AUROC and AUPR are each the area under one of them [40, 41]:

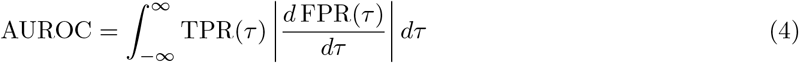

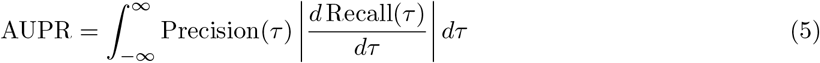

Here *τ* is exactly what separates AUROC/AUPR from accuracy, F1, precision, and recall as usually reported: those four are each evaluated at one fixed *τ*, so a different cutoff gives a different value; change *τ* and accuracy, F1, precision, and recall all change with it. AUROC and AUPR instead integrate over every possible *τ* at once, so both integrals above evaluate to the same number no matter which *τ* a reader might otherwise have picked; *τ* only ever appears as the integration variable, never as an input the final AUROC/AUPR value depends on: the defining property of threshold-independent evaluation as formalized in the ROC-analysis literature [40].

AUROC integrates TPR against FPR. Since *TN* scales with *N*, a fixed *FP* yields a shrinking FPR as *N* grows: for *N ≫ FP*, FPR *≈ FP/N→* 0, so AUROC remains high even as false positives accumulate relative to the small positive class. AUPR instead integrates Precision against Recall (=TPR), and Precision’s denominator *TP* + *FP* scales with *P*, not *N* : the same *FP* that barely moves FPR can dominate *TP* + *FP* when *TP* is small, pulling Precision (and AUPR) down sharply. This asymmetry between AUROC’s insensitivity and AUPR’s sensitivity to the same false positives under *N ≫P* is the formal basis for the relationship between the two curves [41]. It is also the standard justification in the imbalanced-classification literature for preferring AUPR over AUROC in this regime [42], a conclusion echoed outside clinical ML by work on similarly rare-event binary outcomes [43], exactly the regime in which this panel’s worst-case 141.21:1 ratio sits.

A further evaluation-design choice follows from the same reasoning, though it is worth stating precisely rather than as a blanket rule against threshold metrics. AUROC and AUPR, computed by sweeping across every possible threshold rather than committing to one, are this panel’s primary comparison metrics, replacing single-threshold metrics such as accuracy, F1, precision, or recall. This is not a claim that threshold metrics are invalid in general. A model put into clinical use still needs an operating threshold, and its precision/recall or very-major-error/major-error rate at that threshold is what matters operationally at that point [44]. VME/ME is not a hypothetical concern either: both Nguyen et al.’s AMR-GNN [26] and Corvelo Benz et al.’s conformal predictor [10], surveyed in Related Work, report VME/ME directly as part of their own evaluation, underscoring that it is an established, threshold-bound clinical metric in this literature rather than one specific to this panel.

The concern is narrower and specific to comparison under imbalance: a threshold-dependent number reported on its own, without an accompanying threshold-independent metric, can make a mediocre model look strong or a good model look weak purely because of where that one threshold happens to fall relative to the S/R ratio. This is the classic “accuracy paradox,” where a threshold tuned or defaulted toward the majority (S) class inflates accuracy while revealing almost nothing about resistant-isolate detection [45].

Papers that report only threshold-bound metrics under this kind of imbalance therefore risk a misleadingly favorable read of model quality, since the reported number is entangled with an unstated threshold choice rather than reflecting discriminative capacity independent of it. This also motivates treating the “right” operating threshold here as a downstream, context-dependent choice rather than a fixed model property, trading missed-resistance risk (*FN* : predicting S when the isolate is truly R) against unnecessary broad-spectrum escalation (*FP* : predicting R when the isolate is truly S), whose selection depends on whether the intended use is clinical decision support or surveillance [44]. AUROC and AUPR are reported as the metrics that characterize the model’s discriminative capacity independent of that downstream choice, consistent with the standard rationale for threshold-independent evaluation in general classifier comparison [40].

## Results

HetSAGE leads on 4/13 datasets: Cefepime, Piperacillin-T. and Tobramycin in *E*.*coli*, and Fusidic acid in *S*.*aureus*, three of the four sim_raw_ *>* 0.99 cases flagged in Dataset as HetSAGE’s expected regime. MLPClassifier leads or ties on 7/13, including a wide margin on *K*.*pneumoniae* Meropenem, which has the highest imbalance in the panel at 141.21:1 but is also the smallest and noisiest dataset, with std*≈* 0.10–0.17 across all models. GraphSAGE, the homogeneous, raw-only model, trails HetSAGE on most datasets, isolating the biomarker edge, not heterogeneity alone, as the source of HetSAGE’s gains where it does win. LGBM is consistently weakest, reflecting that a single boosting model on the full 6,000-dim raw spectrum, without any graph or consensus structure, is the weakest baseline in this panel.

AUROC tells a starker version of the same story: MLPClassifier leads on 10/13 datasets, HetSAGE and GraphSAGE each lead on 1 dataset, SVM on 1, and LGBM on none. This matches the known AUROC/AUPR divergence under class imbalance: AUROC rewards ranking quality across both classes and is less sensitive to the minority (R) class than AUPR, so it understates exactly where the graph’s biomarker-edge advantage concentrates: the sparse-R, high-imbalance datasets highlighted in Table 5. The AUPR gap for HetSAGE is real but narrower than AUROC alone would suggest a general-purpose model is being beaten by; AUPR remains the more decision-relevant metric for this task, per the asymmetry established in Evaluation Metrics and consistent with Dataset’s framing.

## Discussion

Table 5 above is a post hoc read of Table 3 and Table 4 (reported in Experiments and Results) against each dataset’s similarity and imbalance in Table 1, serving as a capacity check for which dataset regime favors MLP versus HetSAGE rather than a separate deep-learning result of its own; see Dataset and Method for the corresponding motivation and architecture.

**Table 3.**
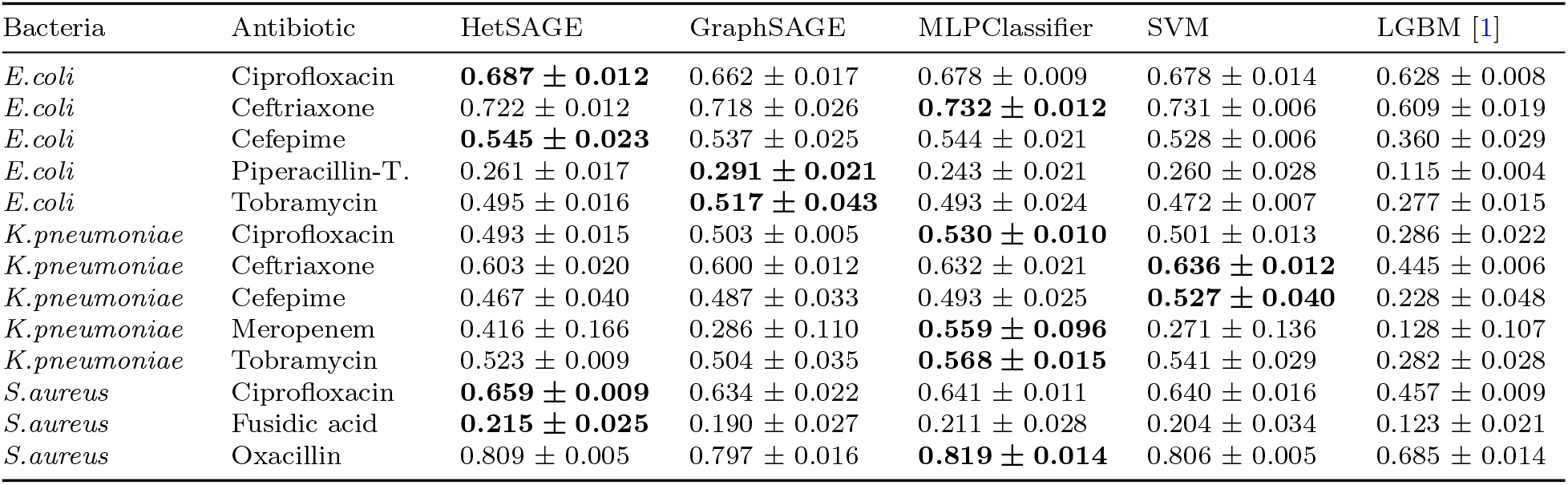
AUPR by Model, All 13 Datasets (mean ± std).

**Table 4.** AUROC by Model, All 13 Datasets (mean ± std).

| Bacteria | Antibiotic | HetSAGE | GraphSAGE | MLPClassifier | SVM | LGBM [1] |
| --- | --- | --- | --- | --- | --- | --- |
| *E.coli* | Ciprofloxacin | 0.812 ± 0.009 | 0.810 ± 0.004 | **0.823 ± 0.004** | 0.816 ± 0.004 | 0.787 ± 0.005 |
| *E.coli* | Ceftriaxone | 0.865 ± 0.004 | 0.857 ± 0.009 | **0.867 ± 0.006** | 0.858 ± 0.006 | 0.806 ± 0.009 |
| *E.coli* | Cefepime | 0.856 ± 0.020 | 0.861 ± 0.011 | **0.868 ± 0.017** | 0.856 ± 0.018 | 0.810 ± 0.019 |
| *E.coli* | Piperacillin-T. | 0.692 ± 0.026 | 0.696 ± 0.016 | 0.685 ± 0.023 | **0.698 ± 0.015** | 0.613 ± 0.019 |
| *E.coli* | Tobramycin | 0.797 ± 0.006 | 0.806 ± 0.024 | **0.818 ± 0.011** | 0.815 ± 0.010 | 0.699 ± 0.005 |
| *K.pneumoniae* | Ciprofloxacin | 0.782 ± 0.015 | 0.782 ± 0.004 | **0.802 ± 0.006** | 0.788 ± 0.012 | 0.701 ± 0.006 |
| *K.pneumoniae* | Ceftriaxone | 0.801 ± 0.013 | 0.791 ± 0.014 | **0.821 ± 0.013** | 0.816 ± 0.007 | 0.750 ± 0.005 |
| *K.pneumoniae* | Cefepime | 0.824 ± 0.023 | 0.800 ± 0.022 | **0.830 ± 0.016** | 0.814 ± 0.015 | 0.737 ± 0.009 |
| *K.pneumoniae* | Meropenem | 0.903 ± 0.082 | 0.866 ± 0.069 | **0.961 ± 0.045** | 0.868 ± 0.052 | 0.661 ± 0.148 |
| *K.pneumoniae* | Tobramycin | 0.801 ± 0.012 | 0.805 ± 0.010 | **0.841 ± 0.022** | 0.827 ± 0.020 | 0.686 ± 0.012 |
| *S.aureus* | Ciprofloxacin | **0.857 ± 0.009** | 0.845 ± 0.009 | 0.849 ± 0.005 | 0.854 ± 0.013 | 0.786 ± 0.011 |
| *S.aureus* | Fusidic acid | 0.687 ± 0.037 | **0.706 ± 0.028** | 0.692 ± 0.024 | 0.679 ± 0.030 | 0.639 ± 0.039 |
| *S.aureus* | Oxacillin | 0.912 ± 0.004 | 0.908 ± 0.009 | **0.917 ± 0.010** | 0.912 ± 0.005 | 0.845 ± 0.012 |

**Table 5.** HetSAGE vs. MLP: Similarity Judged First, Imbalance Second.

| Order | Threshold | Empirical Pattern | Predictive Strength (This Panel) |
| --- | --- | --- | --- |
| 1st | $\text{sim}_{\text{raw}} > 0.99$ (high similarity) | Graph structure (HetSAGE or GraphSAGE) wins | Strong: 4/4 high-similarity datasets (100%) vs. 2/9 otherwise (22%) |
| 2nd | Imbalance ratio $> 10$ (high imbalance) | Graph structure wins somewhat more often | Weak: 3/5 high-imbalance datasets (60%) vs. 3/8 otherwise (37.5%) |

The two thresholds are not equally predictive: spectral similarity is checked first because it is the dominant, near-total split in this panel, while imbalance ratio is a secondary, much weaker signal that only nudges the odds and does not override similarity. The one dataset where the two disagree is *E.coli* /Ciprofloxacin (ratio 2.95:1, below the low-imbalance threshold, but sim_raw_ 0.9931, above the high-similarity threshold); it resolves in favor of similarity: HetSAGE wins on AUPR (0.687 vs. MLP’s 0.678) despite the low imbalance that would otherwise favor MLP. This condition-by-condition split is an empirical pattern observed across the 13 datasets in this panel, not a general rule derived from a controlled sweep over similarity or imbalance; confirming it would require testing additional datasets spanning these same regimes.

## Conclusion

This panel’s evaluation shows that HetSAGE’s advantage over a plain MLP is conditional rather than universal, concentrated in the high-spectral-similarity, high-imbalance regime identified above, where a dedicated biomarker edge and per-edge-type dropout give the graph aggregator a genuinely informative second view rather than a second noisy one. Outside that regime, a well-tuned MLP remains competitive or better, so the practical guidance from this panel is to check a dataset’s similarity and imbalance profile before choosing the heavier heterogeneous-graph architecture over a simpler baseline.

One direction remains open beyond the two data-driven edge types validated here (raw, biomarker). This panel currently trains its 13 species-antibiotic datasets independently; whether species- or antibiotic-level relatedness or grouping, such as taxonomic proximity or shared antibiotic class/mechanism-of-action, could serve as an additional graph view or a shared-parameter signal across panels remains an open question. Unlike a naive isolate-linking edge, grouping at the species/antibiotic level uses only known experimental metadata, not inferred genotype, so it carries no circularity risk.

## Supporting Information

**S1 Appendix. Feature Selection Comparison Figures**.

Referenced from Consensus-Vote Biomarker Construction as supporting evidence that the biomarker consensus vote is not a black box; reproduced here rather than inline since the main text only needs to point to them, not walk through them in place. These two datasets, not all 13 in the panel, are shown because *K*.*pneumoniae*/Meropenem and *S*.*aureus*/Oxacillin are the only pairs for which an independently published clinical or ML-literature reference marker exists to compare the consensus-vote peaks against; the remaining 11 datasets have no equivalent external marker in the literature, so this comparison is illustrative rather than an exhaustive validation across the full panel.

Fig 5 shows this comparison for *K*.*pneumoniae*, Meropenem: LR, RF, LGBM, and Mann-Whitney each independently select their own candidate m/z peaks (top four rows), and the 3/4 (red) and 4/4 (gold) consensus bands cluster tightly rather than scattering randomly across the 2,000–20,000 Da range: visual confirmation that the vote is converging on a small, consistent set of peaks rather than four disagreeing lists. The 3/4-consensus peak at m/z 11,399 sits adjacent to the clinical (pKpQIL, green: same p019/11,109 Da marker family cited above) [35–37] reference band, matching the value reported in the main text; the 4/4-consensus peaks at m/z 3,029 and 4,586 sit adjacent to the ML-literature (LGBM, blue) [38] reference band.

Fig 6 shows the same convergence pattern in *S. aureus* Oxacillin: the four methods again cluster into tight 3/4 and 4/4 consensus bands rather than producing disagreeing lists, and the 4/4 consensus peaks at m/z 2,852 and 6,011, which land near the clinical PSM-mec/delta-toxin (green) [33, 34] and ML-literature (blue) [39] reference bands: the same PSM-mec proximity referenced numerically above.

## Data Availability

The DRIAMS dataset (DRIAMS-A, B, C, and D) used in this study is publicly available via Dryad at https://datadryad.org/stash/dataset/doi:10.5061/dryad.bzkh1899q, as described in [1]. No new patient data were collected. Predicted resistance probability scores for all five baseline classifiers across 13 bacterium-antibiotic combinations and three random seeds are provided in Supplementary Data S1.

